# The adoptive transfer of BCG-induced T lymphocytes contributes to hippocampal cell proliferation and tempers anxiety-like behavior in immune deficient mice

**DOI:** 10.1101/844704

**Authors:** Dan Song, Fangfang Qi, ShuaiShuai Liu, Zhongsheng Tang, Jinhai Duan, Zhibin Yao

## Abstract

We previously have reported that neonatal Bacillus Calmette-Guerin (BCG) vaccination improves neurogenesis and behavior in early life through affecting the neuroimmune milieu in the brain, but it is uncertain whether activation phenotypes and functional changes in T lymphocytes shape brain development. Here, we studied the effects of BCG vaccination via the adoptive transfer of T lymphocytes from the BALB/c wild-type mice into naive mice. Our results show that mice adoptive BCG-induced lymphocytes (BCG->naive mice) showed anxiolytic and antidepressant-like performance when completing an elevated plus maze (EPM) test. Meanwhile, BCG->naive mice possess more cell proliferation and newborn neurons than PBS->naive and nude mice in the hippocampus. IFN-γ and IL-4 levels in the serum of BCG->naive mice also increased, while TNF-α and IL-1β levels were reduced relative to those of PBS->naive and nude mice. We further found that BCG->naive mice showed different repartition of CD4^+^ and CD8^+^ T cell to naive (CD62L^+^ CD44^low^), effector memory (CD62L^−^ CD44^hi^), central memory (CD62L^+^ CD44^hi^) and acute/activated effector (CD62L^−^ CD44^low^) cells in the spleen. Importantly, the adoptive transfer of BCG-induced T lymphocytes infiltrated into the dura mater and brain parenchyma of the nude mice. Activation phenotypes and functional changes in T lymphocytes are very likely to affect the neuroimmune milieu in the brain, and alterations in ratios of splenic CD4^+^ and CD8^+^ memory T cells may affect the expression of correlative cytokines in the serum, accounting for our behavioral results. We conclude thus that the adoptive transfer of BCG-induced T lymphocytes contributes to hippocampal cell proliferation and tempers anxiety-like behavior in immune deficient mice. Our work shows that BCG vaccination improves hippocampal cell proliferation outcomes and behaviors, likely as a result of splenic effector/memory T lymphocytes regulating the neuroimmune niche in the brain.

## 2. Introduction

Bacillus Calmette - Guerin (BCG) is listed as an immunization vaccine for infants and young children in most countries. We have reported in previous works that neonatal BCG vaccination improves the outcomes of neurogenesis and the behaviors of young mice and this effect may be associated with a systemic Th1 bias [1]. At the same time, accumulating evidence has shown that T lymphocytes modulate behavior, cognition and adult hippocampal neurogenesis through direct interactions with the central nervous system (CNS). These effects are mostly mediated by helper T lymphocytes, a subset of CD4^+^ T lymphocytes differentiated by CD3^+^ T lymphocytes [2–4].

It is widely known that there is a reciprocal association between the CNS and peripheral circulation and especially within the immune system, but whether BCG vaccination affects brain functions by altering the activation phenotypes and functions of T lymphocytes. There is no direct evidence. To address this, we designed an experiment in which T lymphocytes drawn from BCG- or PBS-injected mice were transferred to lymphopenic nude mice. The stain matched nude mice repopulated with T lymphocytes cells drawn from BCG-injected mice exhibited less anxious and depressive-like behaviors and increased levels of hippocampal cell proliferation than the PBS-injected control mice. Studies have shown that anatomical structures of the meninges such as the choroid membranes/plexus (ChP) connect the peripheral blood system to the brain via CSF circulation [5, 6]. We thus also examined whether T lymphocytes are recruited to the meninges or are even infiltrated to the brain parenchyma through ChP. We found that CD3^+^ lymphoid cells of BCG->naive mice densely distribute in the meninges across all levels of the brain and especially in the ChP. Our study further shows that altered activation status of T lymphocytes may affect the expression of related cytokines, such as IFN-γ, TNF-a, IL-4 and so on, which may account for the neuroprotective role of neonatal BCG vaccination.

## 3. Materials and methods

### 3.1 Animals and housing conditions

All procedures performed were approved of by the Animal Care and Use Committee of Sun Yat-sen University and conformed to the Guide for the Care and Use of Laboratory Animals of the National Institutes of Health, USA. BALB/c mice (6 weeks old) used as donors of lymphocytes and 6 weeks old BALB/c background nude mice used as recipients of lymphocytes were purchased from the Sun Yat-sen University Laboratory Animal Center (Guangzhou, China). The choice of BALB/c mice as donors is based on the background genotype of the nude recipient mice, although BALB/c mice are known to have a Th2 bias. Wide-type mice used across all experiments are also BALB/c mice. Both the BALB/c mice and the immune deficient nude recipients were male mice and housed in a specific pathogen-free facility. The colony was maintained on 12-h light/dark cycles with free access to food and water in a temperature- and humidity-controlled room.

### 3.2 Vaccination

The immune procedure of BCG used (D2-BP302 strain, Shanghai Institute of Biological Products, Shanghai, China) according to our previous study with slight modifications [1]. Each mouse was injected with 100 μl BCG suspension containing 4×10^5^ CFU (i. h.) or equally sterile PBS for once at 6-week old.

### 3.3 Adoptive transfer

After 7 days of BCG injection, the spleens of donor mice were removed and homogenized by syringe plunger into a single-cell suspension using RPMI Medium 1640 basic (Life Technologies, Beijing, China) supplemented with 10% fetal bovine serum and antibiotics. Red blood cells were lysed with ACK Lysis Buffer (1X, 555899, Biolegend). Splenocytes were washed and resuspended in isolation buffer at a concentration of 1×10^8^ cells/ml. For CD3^+^ isolation, splenocytes were further separated by negative selection using the EasySep T cell isolation kit designed for CD3^+^ T cells (Stem Cell Technologies, Vancouver, BC, Canada) following the manufacturer’s guidelines for manual separation. 8 donors per recipient in the present study, and separated CD3^+^ cells were pooled for transfers. Each nude mouse was reconstituted with 10 million cells with a total volume of 200 µl via tail vein injection. Recipient nude mouse received CD3^+^ T cells from a BCG-injected mouse were defined as BCG->naive. PBS->nude mouse was generated via adoptive transfer of CD3^+^ spleen cells from a PBS-injected mouse into a naïve BALB/c background nude mouse. Naive was defined by injection of a total volume of 200 µl PBS via tail vein into a BALB/c background nude mouse. Normal BALB/c mice are considered as the wide-type group (WT). WT, BCG->naive, PBS->naive and Naive mice were behaviorally phenotyped through open-field and elevated plus maze tests 14 days after T cell injection.

### 3.4 Behavioral Test

#### 3.4.1 Open-field test (OFT)

For open-field exploration, mice were placed in a corner of a dimly lit chamber of the open-field arena 14 days after injection, and the time spent in the peripheral and center zone were monitored using the TopScanTM 2.0 system (Clever Sys. Inc). Mouse locomotor activity was traced and quantified during 5 min as described previously [3]. After each trail, 70% ethanol was used to thoroughly clean the apparatus.

#### 3.4.2 Elevated plus maze (EPM)

The mice were subjected to the EPM task one hour after the OFT task was completed on the same day. The mice were individually placed in the central area facing a constant open arm and were allowed to freely explore the apparatus for 5 min. The behavior of the animals was recorded with a video tracking system (Noldus Etho Vision XT, the Netherlands). The amount of time spent in the open arm, the number of entries made into the open arm and the number of times mice arrived at the end points of the open arms were analyzed. The apparatus was cleaned using 70% ethanol after each mouse completed the test as in the OFT.

### 3.5 Flow cytometry

After behavioral tests were conducted, spleens from different groups were harvested and mechanically dissociated using a syringe. Single cell suspension was filtered through a 70 μm nylon mesh. Next, cells were assayed for surface antigens by flow cytometry. In brief, leukocyte cells were incubated with rat anti-mouse CD3-PE (1.25μl/Test, 554833, Biolegend), CD4-PE-Cy5 (0.5μl/Test, 553050, Biolegend), CD8a-FITC (0.5μl/Test, 553031, Biolegend), CD44-APC (0.5μl/Test, 559250, Biolegend) and CD62L-APC-A750 (0.5μl/Test, AB_10392694, eBioscience) antibodies for 40 minutes in 4 °C. For intracellular IFN-r (0.5μl/Test, AB_1272169, eBioscience) and IL-4 (0.5μl/Test, AB_10714980, eBioscience) detection, leukocyte single cells were cultured with DMEM (HyClone) supplemented with 10% FCS, 1 mm L-glutamine, 100 U/ml penicillin, 100 mg/ml streptomycin (1:1000; BD Biosciences) and for 2 h at 37 °h and incubated with PMA (10 ng/ml; Sigma-Aldrich) ionomycin (250 ng/ml; Sigma-Aldrich) and Brefeldin-A (10 ug/ml) for 5 h, cells were washed, fixed, permeabilized using Fixation/Permeabilization Kit (1X, 4338045, BD Biosciences) according to the protocol of the manufacturer. Cells were analyzed on a CytoFLEX S (Beckman Coulter) after antibodies were washed.

### 3.6 Immunofluorescence

After harvesting the spleens, the mice were perfused with cold PBS to remove peripheral blood, after removal of posterior cervical muscle, snip the skull from foramen magnum to maxillae, the top of the skull was removed with surgical scissors. The brains of the mice were excised from the skull and fixed overnight in 4 % paraformaldehyde at 4 °C for 12 h and then dehydrated with 10%~30% sucrose at 4 °C for another 72 h. The brains were then sliced into serial coronal sections (40 µm) on a freezing microtome (Leica SM2000R; Leica Microsystems GmBH, Wetzlar, Germany). For cell proliferation analysis, the mice were given three BrdU (B5002, Sigma-Aldrich, St. Louis, MO, USA) injections (50 mg/kg, i. p., once every 2 h) prior to excising. Meninges attached to the skull cap were fixed in 2 % paraformaldehyde (PFA) for less than 24 h at 4 °C. The dura was then peeled with microscissors from the skullcap, rinse in PBS buffer for 3 times, and then whole meninges were placed on polylysine-coated slides for immunofluorescence assay.

The following primary antibodies were used: anti-BrdU monoclonal antibodies (1:400, OBT0030, Oxford Biotechnology, UK), goat anti-DCX (1:400, sc-8066, Santa Cruz Biotechnology, CA, USA), rat anti-mouse anti-CD3 (1:400, 14-0032-82, eBioscience, Santa Clara, CA, USA), and rabbit anti-CD4 (1:400; 210846, Sigma-Aldrich, St. Louis, MO, USA). The following secondary antibodies were used: Alexa Fluor 594 donkey anti-rat, Alexa Fluor 488 donkey anti-goat, Alexa Fluor 488 goat anti-rabbit, Alexa Fluor 555 goat anti-rat, and Alexa Fluor 488 goat anti-mouse antibodies (1:1000; all from Invitrogen, Carlsbad, CA, USA).

### 3.7 Photography and confocal imaging

Image acquisition and analysis were systematically performed using a confocal microscope (LSM 780, Carl Zeiss, Germany) with 20 × and 40 × objectives. Quantitative analyses of BrdU^+^ and DCX^+^ cells in the DG were completed using the optical-fractionator method with a stereology system (Stereo investigator, Micro Bright Field, Inc., Williston, VT, USA) as described previously [3]. The anterior 50 % of the structure was defined as the dorsal hippocampus. The number of total dorsal BrdU^+^, DCX^+^ cells were measured in an equidistant series of six coronal sections (240 μm apart) using the stereology system.

### 3.8 Enzyme-linked immunosorbent assay (ELISA)

Concentrations of IFN-γ, IL-4, TNF-a, and IL-1β in the serum were measured as described previously [4]. In brief, blood samples were collected from all groups in centrifuge tubes and incubated at room temperature for 2 h, followed by an overnight incubation at 4 °C before the serum was isolated. Blood samples were centrifuged at 4 °C (4,000 rpm for 15 min), and sera were transferred to another set of tubes. The sera and groups were stored at −80 °C until use. Standard curves were prepared by serially diluting the standard with the kit assay buffer in concentrations ranging from 1000 pg/ml to 7.8 pg/ml.

## 4. Statistics

The results were analyzed using one-way ANOVA with the treatment WT, BCG->naive, PBS->naive and naive mice used as independent variables. When warranted, post hoc LSD tests for multiple comparisons and planned contrast analyses (Student’s *t* tests) were performed. All data are presented as the means ± SEM. *p* values of < 0.05 were considered statistically significant.

## 5. Results

### 5.1 Effect of the adoptive transfer of T lymphocytes on anxious-like behavior

Mice were subjected to an OFT and EPM test 14 days after adoptive transfer of T lymphocytes. For the EPM test for anxiety, the BCG->naive mice spent more time in the open arms than the PBS->naive and nude mice (Fig. 1A; *F* _(3,31)_ = 13.137, \**p* < 0.05) and spent considerably more time in the end of open arms (Fig. 1C; *F* _(3,32)_ = 4.890, \*\**p* < 0.01; *n* = 5-12), denoting an anxiolytic effect of BCG vaccination. From the OFT test we found that BCG->naive mice tended to spend more time in center zone than PBS->naive and nude mice, but these results are not statistically significant (Fig. 1F, *F* _(3,27)_ = 0.793, *p* = 0.508; *n* = 6-10). These findings suggest that the reconstitution of nude mice with BCG-induced lymphocytes tends to temper anxiety-like behavior.

**Fig. 1.**
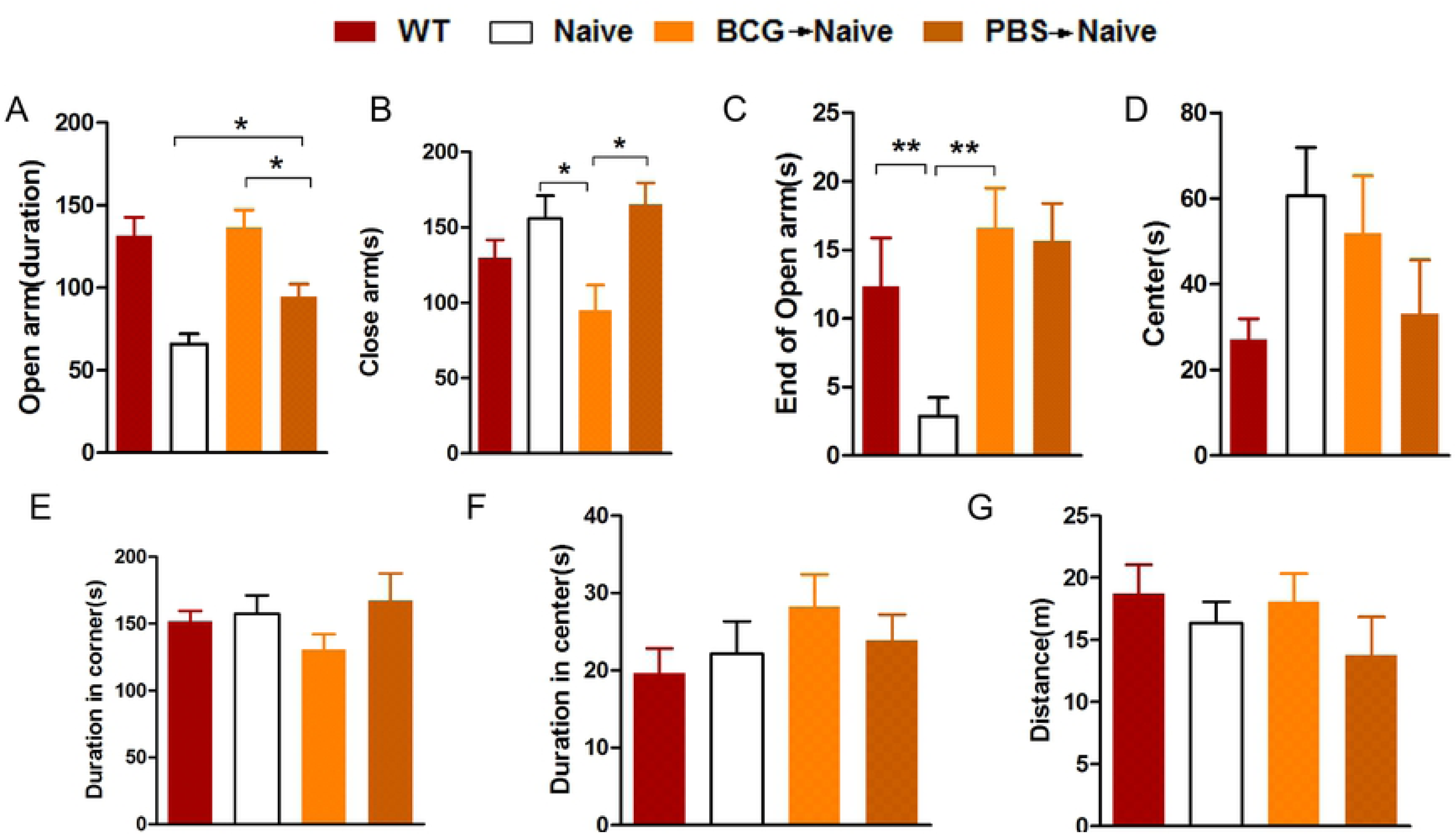
The BCG->naive mice showed reduced levels of anxiety-like behavior in the EPM and OFT. Bars represent the average amount of time spent in open arms, in close arms, at the ends of open arms and at the center of each group of mice for the EPM (A-D). Bars represent the average amount of time spent in corners and central areas and the distance travelled by each group of mice for the OFT (E-G). Data represent means ± SEM. \**p* < 0.05, \*\**p* < 0.01; *n* = 5-12/group (because the nude mice have low immunity and are prone to death, the number of mice in the study ranges widely).

### 5.2 The adoptive transfer of BCG-induced T lymphocytes promotes hippocampal cell proliferation

Studies have shown that immune system affect CNS functioning and structures via humoral and cellular pathways conveying signals to the brain [2]. As nuclear BrdU labels newborn cells, DCX is a specific marker of newborn neurons. We thus assessed whether BCG-induced T cells promote hippocampal cell proliferation in the DG. In line with previous research [7], in immune-deficient nude mice, the hippocampal cell proliferation was markedly impaired, and could be restored by adoptive transfer of PBS-induced T lymphocytes. Notably, the data show that BCG->naive mice have significantly more BrdU^+^ and DCX^+^ cells than both PBS->naive and nude mice (Fig. 2; BrdU^+^, *F* _(3,20)_ = 26.354, \*\*\**p* < 0.001; DCX^+^, *F* _(3,20)_ = 26.059, \*\*\**p* < 0.001; *n* = 6). These results show that BCG-induced T lymphocytes contribute to hippocampal cell proliferation and temper anxiety-like behaviors, suggesting that BCG-induced T lymphocytes promote hippocampal cell proliferation at least in part through immune profiles of the T lymphocytes induced by BCG.

**Fig. 2.**
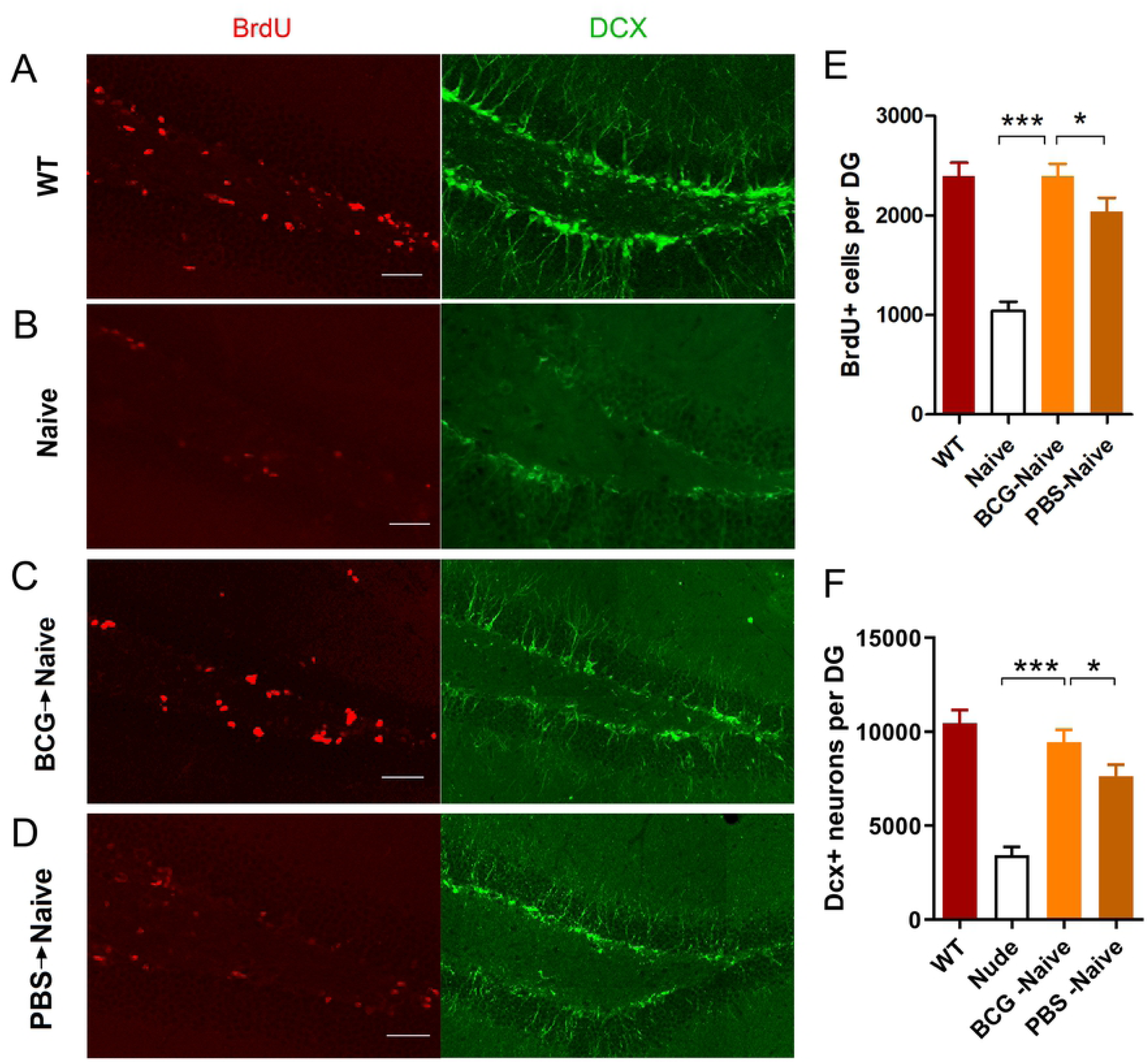
The BCG->naive mice showed increased levels of hippocampal neurogenesis (A-F). Representative confocal micrographs of BrdU^+^- and Dcx^+^-labeled cells of the DG for each group of mice (A-D). Bars represent average values (mean ± SEM) for BrdU^+^- or Dcx^+^-labeled cells of each group (E-F). \**p* < 0.05; \*\*\**p* < 0.001; *n* = 6/group. Scale bar: 50 μm in A-D.

### 5.3 BCG vaccination alter the activation status of peripheral T lymphocytes

Studies have shown that CD4^+^ T lymphocyte activities provide a neuroimmunological link in the control of adult hippocampal neurogenesis [6, 8]. We thus explored whether increased levels of hippocampal cell proliferation are related to the differentiation of T lymphocytes to CD4^+^ cells. After specific CD3^+^ aggregation, purity detection results show a proportion of adoptive CD3^+^ T cells of more than 97%. Fourteen days after adoptive transfer, single cells taken from the spleens of the BCG->naive, PBS->naive and nude mice were incubated with CD3, CD4, CD8, CD44 and CD62L antibodies for flow cytometry analysis. We found no differences in CD3^+^ T cell populations both in spleen and peripheral blood between BCG->naïve and PBS->naive mice (Fig. 3A, B, E, F. BCG->naïve vs. PBS->naïve: *p* > 0.05). However, BCG->naive mice present much higher ratios of CD4^+^ T cells to total lymphocytes both in spleen and peripheral blood (Fig. 3C, G. Spleen; *F* _(3, 20)_ = 264.114, \*\*\**p* < 0.05; Fig. 3D, H. peripheral blood: *F* _(3, 20)_ = 101.124, \*\*\**p* < 0.05; *n* = 6). Interestingly, no significant differences in the percentage of CD3^+^, CD4^+^ and CD8^+^ T cells were observed between WT mice and BCG-vaccinated mice (Fig. 4D-F; CD3: *p* = 0.058; CD4: *p* = 0.668; CD8: *p* = 0.462; Student *t* test; *n* = 7).

**Fig. 3.**
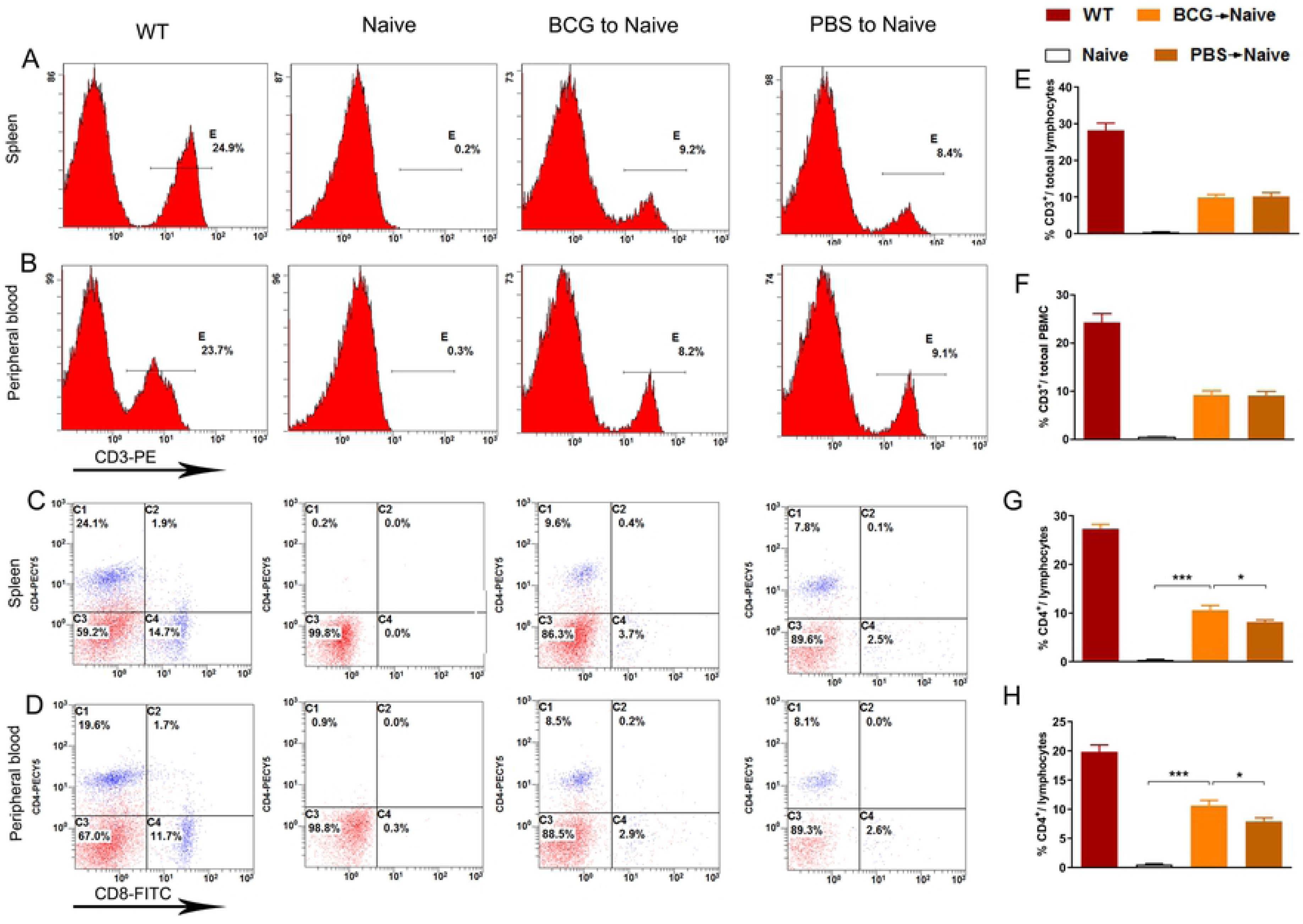
The BCG->naive mice showed more CD4^+^ in spleen and peripheral blood (A-H). FACS: the proportion of CD3^+^ cells in spleen and peripheral blood for each group (A-B); the proportion of CD4^+^ and CD8^+^ cells in spleen and peripheral blood for each group (C-D); bars denote the proportion of CD3^+^ and CD4^+^ to total lymphocytes found in the spleen and peripheral blood of each group (E-H). \**p* < 0.05, \*\**p* < 0.01; *n* = 6/group.

**Fig. 4.**
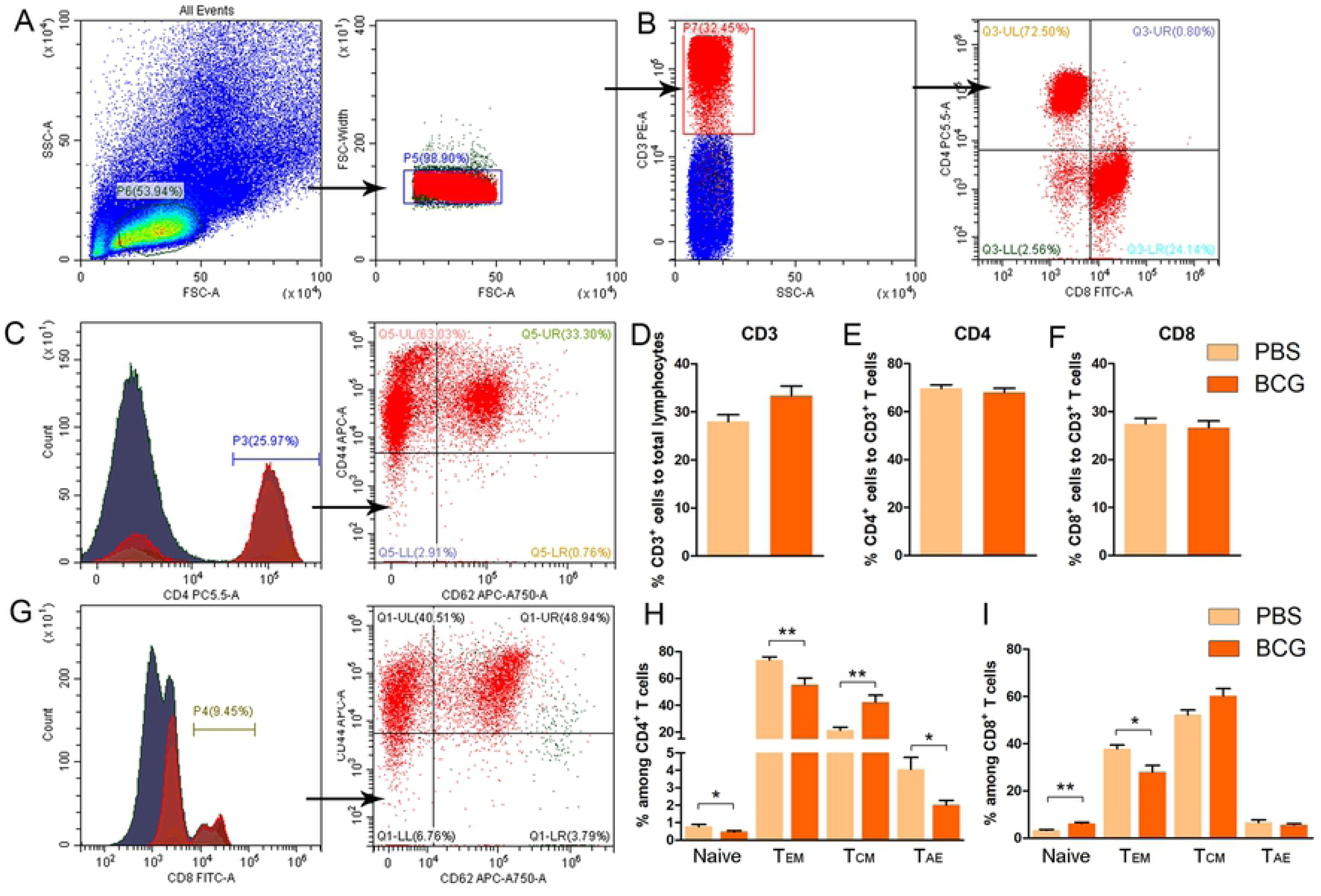
The differentiation of CD3^+^ T lymphoid cells in wild type mice after BCG vaccination (A-I). No significant differences in the percentage of CD3^+^, CD4^+^ and CD8^+^ T cells were observed in WT mice after BCG vaccination (A, B, D-F). Different activation status of CD4^+^ and CD8^+^ T cells in the spleen of WT mice after BCG vaccination. (C, G, H, I). Data represent means ± SEM. \**p* < 0.05, \*\**p* < 0.01; *n* = 6-7/group.

In parallel, we also evaluated the activation status of CD4^+^ and CD8^+^ T cells in the spleen of WT mice vaccinated BCG or not based on expression of CD44 and CD62L activation markers. Although BCG vaccination has no effects on ratios of CD8^+^ or CD4^+^ to CD3^+^ in the spleens of WT mice, the data show that the frequency of naive (CD62L^+^ CD44^low^) CD8^+^ T cells is significant increased (Fig. 4I. Naïve: \*\**p* < 0.01; Student *t* test; *n* =6-7); whereas the counts of effector memory (CD62L^−^ CD44^hi^) CD8^+^ T cells is obviously decreased in BCG-vaccinated wide-type mice as compared to the control mice that received PBS (Fig. 4I. Tem: \**p* < 0.05; Student *t* test; *n* =6-7). No significant differences were observed in the frequency of central memory (CD62L^+^ CD44^hi^) or acute/activated effector (CD62L^−^ CD44^low^) CD8^+^ T cells (Fig. 4I. *p* > 0.05 for both). However, we found that naive (CD62L^+^ CD44^low^) or effector memory (CD62L^−^ CD44^hi^) or acute/activated effector (CD62L^−^ CD44^low^) CD4^+^ T cells drop dramatically (Fig. 4H. Naïve: \**p* < 0.05; Tem: \*\**p* < 0.01; Tae: \**p* < 0.05: Student *t* test; *n* =6-7); whereas obvious increase in frequency of central memory (CD62L^+^ CD44^hi^) CD4^+^ T cells was detected in BCG-vaccinated mice as compared to the control mice (Fig. 4H. \*\**p* < 0.01; *n* = 6-7). These results confirmed that different activation status of CD4^+^ and CD8^+^ T cell in the spleen of WT mice after BCG vaccination. Therefore, the immune profile of the donors is different between BCG-induced T cells and PBS-T cells. Interestingly, after the adoptive transfer of T cells to nude mice, the ratio of CD8^+^ to total lymphocytes drops sharply from 14 to 2 percent both in spleen and peripheral blood (Fig. 3C, D). From our experimental results, it can be inferred that increased levels of adult hippocampal cell proliferation in BCG->naive mice might be related to alteration in activation status of adoptively transferred T lymphocytes.

### 5.4 The adoptive transfer of BCG-induced T lymphocytes tempers pro-inflammatory cytokine responses in the periphery

Cytokines and neurotrophins are believed to be significant mediators of the role of peripheral immune activation in brain development and cognitive functioning [5, 9, 10]. To further explore the potential mechanisms of behavioral outcomes, we measured serum cytokine levels. We found levels of IFN-γ and IL-4 significantly increase in the BCG->naive mice relative to those of the PBS->naive and nude mice. Levels of TNF-α and IL-1β were reduced in BCG->naive mice relative to those of the PBS->naive and nude mice (Fig. 5; IFN-γ: *F* _(3, 20)_ = 13.767, \**p* < 0.05; IL-4: *F* _(3, 20)_ = 3.750, * *p* < 0.05; TNF-α, *F* _(3, 20)_ = 2.457, \**p* < 0.05; IL-1β, *F* _(3, 20)_ = 7.072, \**p* < 0.05; *n* = 5-6). These findings show that the adoptive transfer of BCG-vaccinated T lymphocytes tempered pro-inflammatory cytokine expression and enhanced IFN-γ and IL-4 expression, thereby regulating hippocampal neuronal proliferation observed above.

**Fig. 5.**
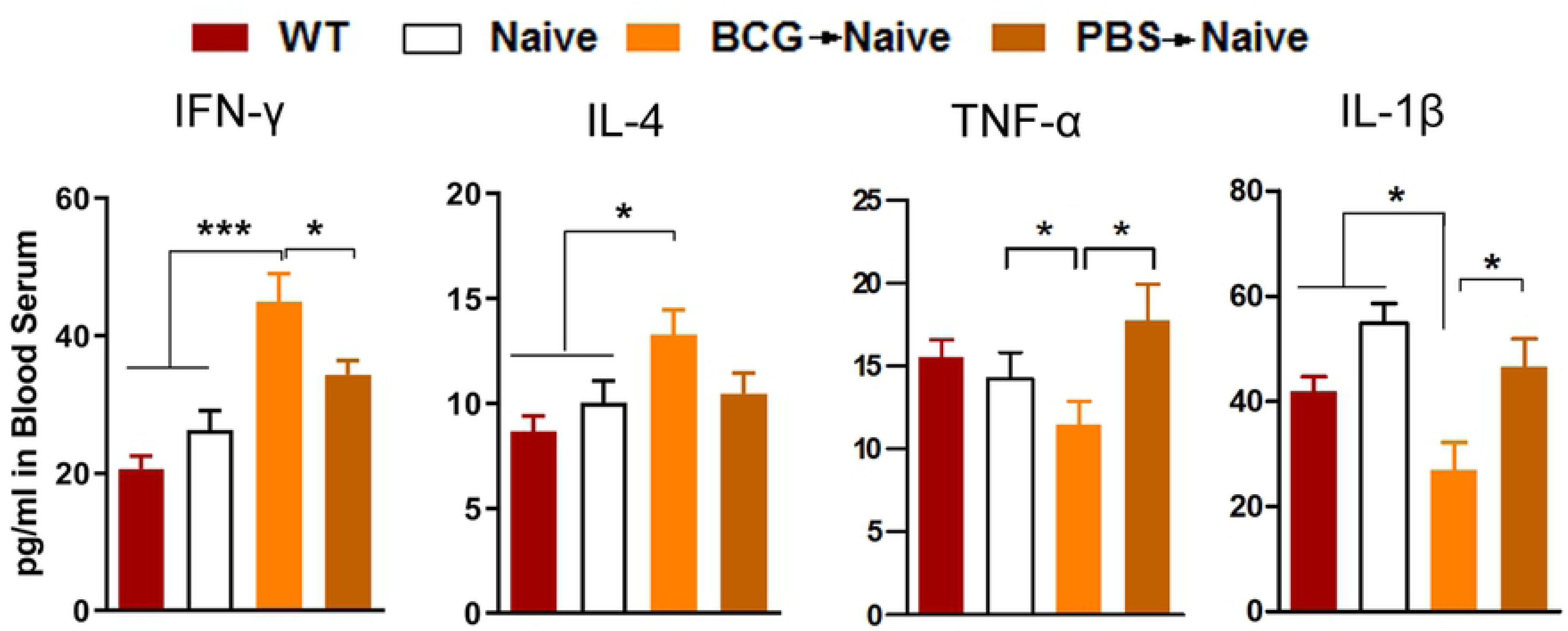
The BCG->naive mice altered serum cytokine levels. Bars denote IFN-γ, IL-4, TNF-α and IL-1β levels for each group. A statistical analysis was performed on primary values of levels between the two groups. Data denote means ± SEM. \**p* < 0.05; \*\*\**p* < 0.001; *n* = 5-6/group.

### 5.5 The infiltration of T lymphocytes in the meninges and hippocampal parenchyma

Hippocampal neurogenesis induced by an enriched environment is associated with the recruitment of T cells to brain parenchyma, which is central to the maintenance of hippocampal neurogenesis and of spatial learning abilities in adulthood [7]. We thus explored whether promoted hippocampal cell proliferation is related to T lymphocyte infiltration in the meninges and brain parenchyma. By immunofluorescence we detected CD3^+^ and CD4^+^ lymphoid cells of BCG->naive mice to be densely distributed in the dura mater with few CD8^+^ lymphoid cells (Fig. 6D, E). CD3^+^ lymphoid cells were found scattered across brain tissues (Fig. 6A) And especially in the choroid plexus (ChP) (Fig. 6B) and ependymal ventriculorum (Fig. 6C) but with less dense areas found in the brain parenchyma close to the ventricle.

**Fig. 6.**
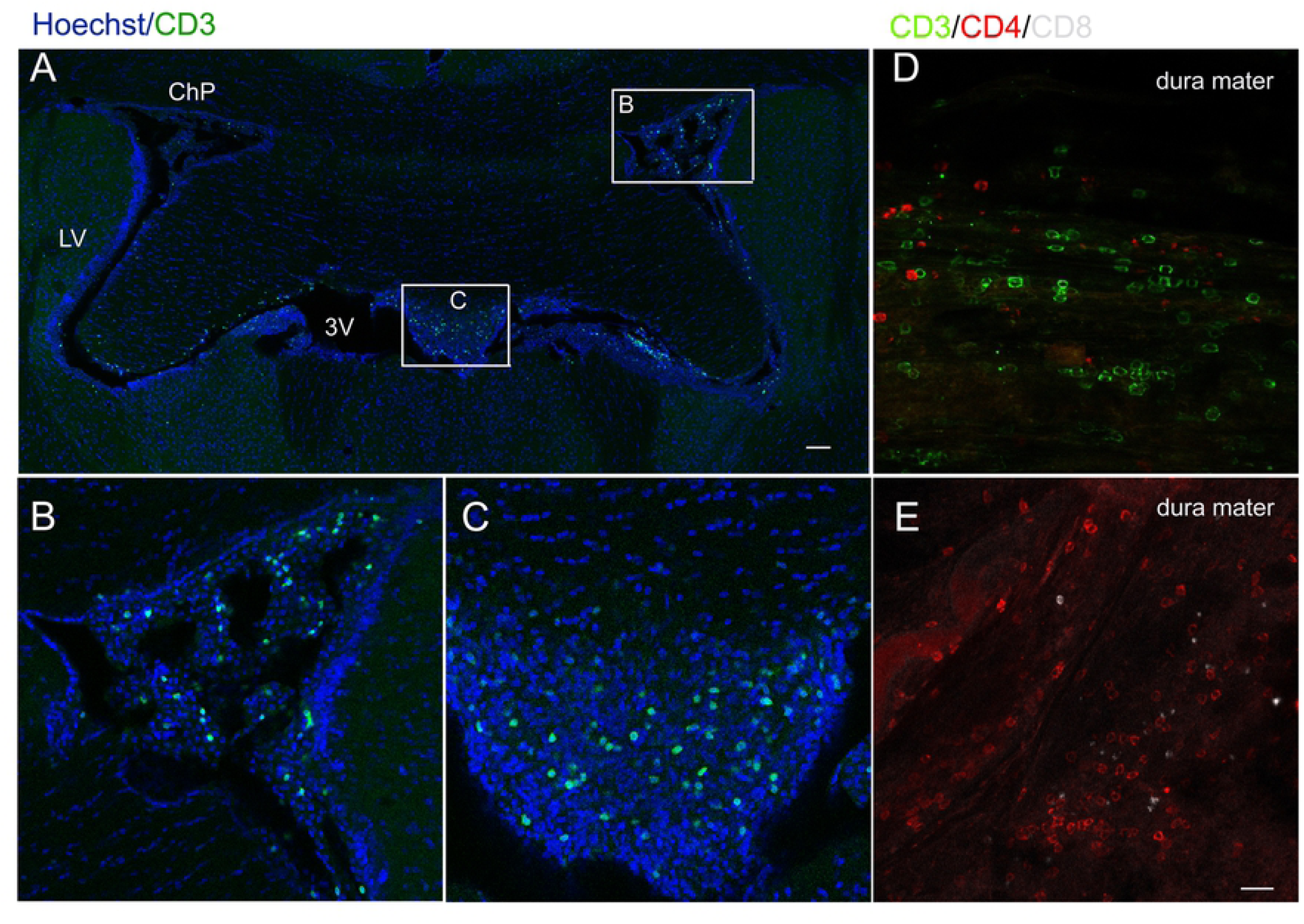
The infiltration of T lymphocytes in the meninges and brain parenchyma after the adoptive transfer of BCG-induced T lymphocytes (A-E). Representative confocal micrographs of CD3^+^ lymphoid cells in the brain slice (A), the choroid plexus (ChP) (B) and the third ventricle (C). Numerous CD3^+^ and CD4^+^ lymphoid cells were found in the dura mater while few CD8^+^ lymphoid cells were found (D, E). Scale bar: 100 μm in A; 20 μm in B-E.

## 6. Discussion

Our results show that the adoptive transfer of T lymphocytes after BCG vaccination contributes to hippocampal cell proliferation and tempers anxiety-like behaviors in nude recipient mice. First, the adoptive transfer of T lymphocytes after BCG vaccination alleviated the anxiety-like behaviors of nude mice and enhanced hippocampal cell proliferation in the DG. Furthermore, we found that the adoptive transfer of BCG-induced T lymphocytes altered immune profiles in vivo while tempering pro-inflammatory cytokine responses in the serum. T lymphocytes permeated through the dura mater and parenchyma close to ventricle were surveyed by immunofluorescence. These parallel alterations are reminiscent of recent results showing that mood states and neurogenesis are typically associated with immune activation characterized by altered inflammatory cytokine expression [11]. Our team has previously shown that neonatal BCG vaccination improves neurogenesis and behavior through affecting the neuroimmune milieu in the brain [1]. However, the function roles of peripheral T lymphocytes in BCG-induced neurogenesis are unclear.

Here, we used immune deficient nude mice without mature T lymphocytes for several sets of experiments. Using adoptive transfer T lymphocytes with different immune properties, we further studied the roles of activation status of T lymphocytes in regulating hippocampal cell proliferation and mood states.

Many studies have shown that specific subsets of lymphocytes can affect hippocampal neurogenesis, spatial learning and memory and stress responses [6, 12–15]. Immune cells (T lymphocytes and microglia) contribute to the maintenance of neurogenesis and of spatial learning abilities in adulthood [10]. At the same time, our previous study shows that the neonatal BCG vaccination of mice improves neurogenesis and behavior via T helper 1 bias [4]. Furthermore, it is now recognized that CD4^+^ T lymphocytes provide a neuroimmunological link in the control of adult hippocampal neurogenesis [6]. From these findings, we speculate that BCG vaccination alters activation status of peripheral T lymphocytes, spurring changes in the secretion of cytokines and neurotrophins in the brain and thus affecting the neuroimmune milieu. Interestingly, FACS analysis of CD4^+^ and CD8^+^ T cells revealed an decrease in the CD62L^−^ CD44^hi^ subpopulation (effector memory T cells) in the spleen of BCG mice compared to the controls, but the opposite was observed in the CD62L^+^ CD44^hi^ CD4^+^ T cells (central memory T cells). Meanwhile, there was an increasing trend in the frequency of central memory (CD62L^+^ CD44^hi^) CD8^+^ T, but these differences were not significant. These results suggested that CD4^+^ and CD8^+^ T cells have different activation phenotypes after BCG vaccination. More recently, a study has revealed that mice raised in enriched environment have different properties in terms of CD8^+^ effector and central memory T cells re-partition in the brain [16]. More importantly, Ritzel et al. [17] demonstrated that a novel population of immunosurveillant CD8^+^ effector memory T cells that represent a hallmark of CNS aging and appear to modify microglia homeostasis under physiological conditions, but are primed to enhance inflammatory response and leukocyte recruitment in pathological conditions. Therefore, it is understandable that the CD4^+^ and CD8^+^ T effector memory T cells are decreased in BCG-vaccinated mice, followed with increased hippocampal cell proliferation in the present study. It will be valuable to determine the roles of memory cells in the hippocampal cell proliferation and anxiety-like behaviors observed in BCG-vaccinated in the future.

Ultimately, BCG-induced T lymphocytes promote hippocampal cell proliferation and temper anxiety-like behaviors in nude mice. From the present study we first show that T lymphocytes can transmit similar functions in their host through preliminary BCG immunization. This result is consistent with recent a report [18] showing that lymphocytes of stressed mice showed less anxiety, increased hippocampal cell proliferation and decreased pro-inflammatory cytokine expression. In brief, Brachman et al. [19] also shows that stress affects no lymphocyte phenotypes prior to transfer. There is a phenomenon considered as a form of behavioral immunization. It means that a moderate-intensity context may immunize the individual, thereby enhancing resistance to the subsequent same stress [18]. So, we conclude that the same underlying mechanism (stressed and immunized mice) should be investigated to explain this consistency in the future.

Song’s group [20] has reported that the brain’s meningeal system and vascular and perivascular spaces serve as homing sites of lymphocytes. Therefore, we also found that the adoptive transfer of T lymphocytes infiltrates into the meninges, choroid plexus and even the brain’s parenchyma. Our recent findings [7] show that the elevated recruitment of T lymphocytes into the CP, dura mater and meningeal inducing meningeal macrophage M2 polarization in response to BCG vaccination serves as evidence of an interior relationship between the periphery and CNS.

In summary, our results provide evidence of effects of BCG-induced T lymphocytes on the modulation of the hippocampal cell proliferation and behaviors of nude mice, which further support the effects of adaptive immune processes on brain and behavior functioning.

## Funding

This work was supported with a grant from the National Natural Science Foundation of China (No. 31700914 and No.31600836), a Special Financial Grant from the China Postdoctoral Science Foundation (2017T100652), a General Financial Grant from the China Postdoctoral Science Foundation (2016M602570) and a Science and Technology Planning Project of Guangdong Province (2017ZC0278).

## Acknowledgments

We thank Dr. Juntao Zou (SYSU), Dr. Kaihua Guo (SYSU), Ms. Qunfang Yuan (SYSU), Dr. Yingying Wu (SYSU), Dr. Yunjie Yang (SYSU) and Dr. Zitian He (SYSU) for their valuable discussions and help with this investigation.

## Competing Interests

The authors have declared that no competing interests exist.

## Author contributions

The study was conceived by Zhibin Yao. Behavior test and Immunofluorescence assays were conducted by Dan Song and Fangfang Qi. FCM was performed by Zhongsheng Tang and Fangfang Qi. Microscopy was performed by Jinhai Duan and Fangfang Qi. The manuscript was written by Fangfang Qi and Zhibin Yao, and edited by Shuaishuai Liu.

